# Looking for a needle in a haystack: magnetotactic bacteria help in “rare biosphere” investigations

**DOI:** 10.1101/2022.07.08.499144

**Authors:** Maria Uzun, Veronika Koziaeva, Marina Dziuba, Lolita Alekseeva, Maria Krutkina, Marina Sukhacheva, Roman Baslerov, Denis Grouzdev

**Affiliations:** Skryabin Institute of Bioengineering Research Center of Biotechnology of the Russian Academy of Sciences, Moscow, Russia; Faculty of Biology, Lomonosov Moscow State University, Moscow, Russia; Department of Microbiology, University of Bayreuth, Bayreuth, Germany; SciBear OU, Tallinn, Estonia

## Abstract

Studying the minor part of the uncultivated microbial majority (“rare biosphere”) is difficult even with modern culture-independent techniques. The enormity of microbial diversity creates particular challenges for investigating low-abundance microbial populations in soils. Strategies for selective sample enrichment to reduce community complexity can aid in studying the rare biosphere. Magnetotactic bacteria, apart from being a minor part of the microbial community, are also found in poorly studied bacterial phyla and certainly belong to a rare biosphere. The presence of intracellular magnetic crystals within magnetotactic bacteria allows for their significant enrichment using magnetic separation techniques for studies using a metagenomic approach. This work investigated the microbial diversity of a black bog soil and its magnetically enriched fraction. The poorly studied phylum representatives in the magnetic fraction were enriched compared to the original soil community. Two new magnetotactic species, *Candidatus* Liberimonas magnetica DUR002 and *Candidatus* Obscuribacterium magneticum DUR003, belonging to different classes of the relatively little-studied phylum *Elusimicrobiota*, were proposed. Their genomes contain clusters of magnetosome genes that differ from the previously described ones by the absence of genes encoding magnetochrome-containing proteins and the presence of unique *Elusimicrobiota*-specific genes, termed *mae*. The predicted obligately fermentative metabolism in DUR002 and lack of flagellar motility in the magnetotactic *Elusimicrobiota* broadens our understanding of the lifestyles of magnetotactic bacteria and raises new questions about the evolutionary advantages of magnetotaxis. The findings presented here increase our understanding of magnetotactic bacteria, soil microbial communities, and the rare biosphere.

## INTRODUCTION

The sequencing of environmental DNA has revealed that the majority of microbial lineages have not been isolated in axenic cultures and have been investigated by culture-independent methods [1, 2], leading to their designation as “microbial dark matter” [3]. In this microbial dark matter, representatives of major phyla occupy the largest part, followed by a “long tail” of rare taxa [4]. Even the use of modern culture-independent techniques sometimes does not allow detailed analysis of the minor components of the uncultured majority, the so-called “rare biosphere” [5]. Identification of rare biosphere representatives, which also play significant roles in biogeochemical cycles, still requires generation of a large amount of metagenomic data [4].

Magnetotactic bacteria (MTB) could help in studies of the rare biosphere due to their magnetic properties [6]. These properties are attributed to the presence of magnetosomes: crystals of magnetite (Fe_3_O_4_) or greigite (Fe_3_S_4_) enveloped by a lipoprotein membrane [7]. Magnetosome synthesis in MTB is controlled by a magnetosome gene cluster (MGC) [8] that comprises the *mam* genes necessary for magnetosome biomineralization and group-specific genes (e.g. *mad*, *man*, *mms*, etc.) that determine the magnetosome shape and size [9]. Bacterial cells use magnetosomes to align along magnetic field lines as they swim using their flagella in a behavior called magnetotaxis [10]. Magnetotaxis is believed to work with both chemotaxis and aerotaxis to assist MTB in maintaining an optimal position at the oxic-anoxic interface in chemically stratified water columns or sediments [11]. These magnetic properties also allow the separation of MTB, a minor component of microbial communities [12], from non-MTB using separation techniques, such as racetrack [13] and MTB trap [14]. MTB of rare biosphere phyla, such as *Riflebacteria*, UBA10199, *Omnitrophota*, and *Fibrobacterota*, were recently detected using these separation techniques [15]. Notably, these bacteria were detected in soils, where MTB have rarely been studied, providing a basis for further study of MTB in this environment.

One rare biosphere phylum is *Elusimicrobiota*. Formerly called “Termite Group 1,” this phylum was first discovered in insect guts [16–18]. A few subsequent studies have increased the number of genomes associated with different habitats, such as groundwater [19], seawater [20], freshwater [21], and wastewater [22]. However, only three *Elusimicrobiota* genomes have been reconstructed from soils [23], perhaps because soils have extremely high microbial diversity, so obtaining rare biosphere genomes is challenging [19]. The first *Elusimicrobiota* genome containing a partial MGC, NORP122, has recently been discovered by genome analysis from the open databases, suggesting that MTB may exist among *Elusimicrobiota* representatives. [24, 25].

Here, we analyzed a microbial community in the magnetically enriched fraction of peaty black bog soil. MTB-CoSe enrichment allowed reconstruction of two novel MTB genomes belonging to two different classes within *Elusimicrobiota*, a comparative analysis of their MGC and predicted metabolic features. Our findings reveal the previously undescribed features of these rare organisms and raise new questions about the potential mechanisms of magnetosome biomineralization. This supplements knowledge about MTB while also providing new information about the *Elusimicrobiota* phylum.

## MATERIAL AND METHODS

### Sampling, microscopy observation, and DNA extraction

Samples of the waterlogged peaty black bog soil were obtained on the slope of the Durykino ravine, Chashnikovo village, Moscow region, Russia (56°02’56.0”N 37°09’53.0”E) in July 2017 [26]. Soil and water samples were collected at a 10 cm depth into plastic containers and transferred to the laboratory for microcosm creation in a 3 L glass bottle. The microcosm was incubated in the dark at room temperature for 15 months, then MTB cells were magnetically enriched using the MTB-CoSe technique [27] by vacuum filtering the soil homogenate through filter paper to remove large soil particles and applying the filtrate to a MiniMACS column (Miltenyi Biotec, Germany) to separate bacteria with magnetic particles, including MTB, from non-magnetic ones. Samples of initial soil homogenate (S) in three replicates, filtrate (F) obtained after vacuum filtration, and magnetically concentrated cells (M) were then used for DNA extraction using DNeasy PowerSoil Kit (Qiagen, The Netherlands). Magnetically enriched cells were observed by transmission electron microscopy (TEM) observations on a JEOL JEM-1011 microscope equipped with an ORIUS SC1000W digital camera and Digital Micrograph (GATAN) software at an 80 kV acceleration voltage. Simultaneously with magnetic enrichment, the physical and chemical parameters of a 50 mL water sample from the microcosm were also determined, and the hydrogen sulfide content was determined in a 10 mL water sample treated with zinc acetate. Anions were analyzed using a Dionex ICS-1100 ion chromatograph (Thermo Scientific, USA) and elements were analyzed by inductively coupled plasma optical emission spectrometry on an Agilent 5110 ICP-OES instrument (Agilent Technologies, USA). Water conductivity was determined with a HANNA HI 2300 conductometric liquid analyzer (Hanna Instruments, USA), water pH and salinity were measured using an Acvilon pH meter (Acvilon, Russia)

### 16S rRNA sequencing, Real-Time PCR

The V3–V4 region of the 16S rRNA amplicons was subjected to high-throughput sequencing using a MiSeq system (Illumina, United States) and MiSeq Reagent Kit v2 (500 cycles) (Illumina, United States), following the manufacturer’s recommendations. The obtained 2×250 bp reads were then processed using workflows implementing the USEARCH v10 scripts [28]. Pair-end reads were demultiplexed (-fastx_demux), merged (-fastq_mergepairs), trimmed to remove the primer sequences (-fastx_truncate), and quality filtered (-fastq_filter). Zero radius operational taxonomic units (zOTUs) were generated by UNOISE3 [29, 30] and assessed using default parameters in the SILVA database (SINA, https://www.arb-silva.de/aligner/, v1.2.11, SILVA reference database release 138.1) [31]. Alpha-diversity of the microbial communities was estimated using the USEARCH v10 -alpha_div command.

The bacterial component in the samples S, F, and M was quantified by real-time PCR using Eub338F/Eub518R primers [32] and SYBR Green I technology in PCR buffer-RB (Syntol, Russia) containing the passive reference dye ROX to normalize the fluorescence signal of the reaction dye. The reaction mix (25 μL) contained 2.5 × PCR buffer-RB (10 μL), each primer (0.25 μL; 20 pM/μL), DNA (5 μL), and MQ water. The CFX96 Touch™ real-time detection system (Bio-Rad, USA) was used for amplification with the following qPCR protocol: polymerase activation 5 min at 95 °C, 10 15 s cycles at 95°C, 45 s at 62°C, 30 15 s cycles at 95°C, and 45 s at 60°C. Samples were analyzed in duplicate and ddH2O (Syntol, Russia) served as a negative control (reaction mix without DNA matrix). Bacterial counts were obtained by comparing the signals from the tested samples with a standard curve prepared by serial dilutions of a standard sample purified using the WizardSV Gel and PCR Clean-Up System Kit (Promega, USA) and subsequently cloned into the pGEM-T vector (Promega, USA) with the target PCR fragment.

### Genome sequencing, assembly, annotation, and metabolic pathway reconstructions

The Genomiphi V2 DNA Amplification Kit (GE Healthcare, USA) was used for whole-genome multiple displacement amplification to generate sufficient DNA for metagenomic sequencing. DNA for metagenomic sequencing was purified by sodium acetate precipitation, and all DNA manipulations used standard protocols. The DNBSEQ (MGI) and Oxford Nanopore Technologies (ONT) platforms were used for short- and long-read DNA metagenomic sequencing, respectively. Short reads were obtained from DNA libraries constructed with the MGIEasy universal DNA library prep set, following the kit protocol. Genomic DNA was sequenced using the DNBSEQ-G400 platform (MGI Tech, China), with 150 bp paired-end reads. FastQC v0.11.9 (http://www.bioinformatics.babraham.ac.uk/projects/fastqc/) was used to assess the read quality, followed by trimming with Trimmomatic v0.39 [33], using the default paired-end read settings.

NEBNext Companion Module for Oxford Nanopore Technologies Ligation Sequencing was used for ONT sequencing library preparation. Sequencing was carried out on a MinION device with a R9.4.1 flow cell (FLO-MIN106D). Guppy v3.4.4 was used to base call, demultiplex, and quality trim the ONT-passed long reads. Long and short trimmed reads were hybrid de novo assembled using SPAdes v3.13.0 with “-meta” flag [34]. Metagenome assembled genomes (MAGs) were reconstructed using MaxBin2 v2.2.7 [35], MetaBAT2 v2.15 [36], and MyCC [37] with standard parameters. Consensus assemblies for the MAGs were chosen using DAS Tool v1.1.3 [38]. Contamination based on taxonomic assignments was removed with RefineM v0.1.2 [21]. The quality metrics were assessed using QUAST v5.0.2 [39]. Genome completeness and contamination were estimated using CheckM v1.2.0 [40]. The taxonomic affiliation of the genomes was determined using GTDB-Tk v2.0.0, release 07-RS207 [41]. Protein-coding sequence identification and primary annotation were performed using the NCBI Prokaryotic Genome Annotation Pipeline (PGAP, v5.3) [42]. The putative MGCs in a set of the *Elusimicrobiota* genomes were screened with MagCluster [43] and further verified using local BLAST and comparison with reference MTB sequences. PFAM and COG domains in MGC protein sequences were detected using the webMGA tool [44].

The metabolic pathways were reconstructed based on the KEGG (Kyoto Encyclopedia of Genes and Genomes) [45] and Distilled and Refined Annotation of Metabolism (DRAM) [46] frameworks. Hierarchical Clustering Analysis (HCA) was used to group the genomes based on the completeness of carbon and energy metabolism pathways estimated by DRAM. Ward’s variance minimization algorithm and Euclidian distance metric were used for clustering. The abundance of genes related to motility was conducted by HCA with the same parameters, except that the numbers of orthologues present were standardized in rows (subtracted from the minimum and divided by maximum) prior to clustering to avoid the bias caused by excessive numbers of certain orthologues present in a genome. Hierarchical clustering was computed in Python using the scipy v1.8.1 library. The resulting dendrogram and heatmap were visualized using the seaborn v0.11.2 library.

### Phylogenetic analyses and genome index calculation

The average nucleotide identity (ANI) values were determined using FastANI v. 1.3 [47]. All *Elusimicrobiota* genomes in NCBI available in March 2022 were selected for genome-based phylogenetic analyses after removal of same-species genomes (ANI > 95%) and genomes with unsuitable GTDB criteria (completeness-5×contamination ≥ 50) [21]. The representative *Elusimicrobiota* genomes from the GTDB r202 database and all known MTB genomes from other phyla were then added to that set. The final set was divided into five groups (groundwater, animal-associated, wastewater, aquatic, and soil) according to NCBI BioSample information.

The GTDB-Tk v.1.7.0 toolkit was used to search for 120 single-copy bacterial marker genes [41]. Maximum-likelihood phylogenetic trees were built with IQ-TREE v1.6.12 [48] using evolutionary models selected by ModelFinder [49]. Branch supports were obtained with 1000 ultrafast bootstraps [50]. Trees were visualized with iTOL v6.5.4 [51]. The 120 single-copy bacterial marker protein-sequence tree (hereinafter called «species tree») was rooted to *Fusobacteriota* [52]. Average amino acid identity (AAI) was calculated using CompareM v0.1.2 [53].

## RESULTS

### Taxonomic structure and abundance of microbial communities before and after magnetic enrichment

The Durykino ravine is located near cultivated fields with soddy-podzolic soil [26]. A stream flowing along the bottom of the ravine causes soil waterlogging. At the sampling time, the air temperature was +25 °C, the sample pH was 6.5, and the salinity was 1%. Part of the formed microcosm of the sampled waterlogged soil was subjected to physical and chemical analysis before the magnetic enrichment (Supplementary Table S1).

Raw sequences of the V3-V4 region 16S rRNA gene were filtered and grouped in zOTUs (Supplementary Table S2). The microbial community composition was analyzed in the initial microcosm soil sample (S), after filtration (F), and after magnetic column enrichment (M). Structural changes in the microbial communities were demonstrated at each stage of separation. The (S) community at the phylum level mostly consisted of *Pseudomonadota* (21.3%), *Thermodesulfobacteriota* (11.4%), and *Acidobacteriota* (10.2%) (Fig. 1A).

**Fig. 1:**
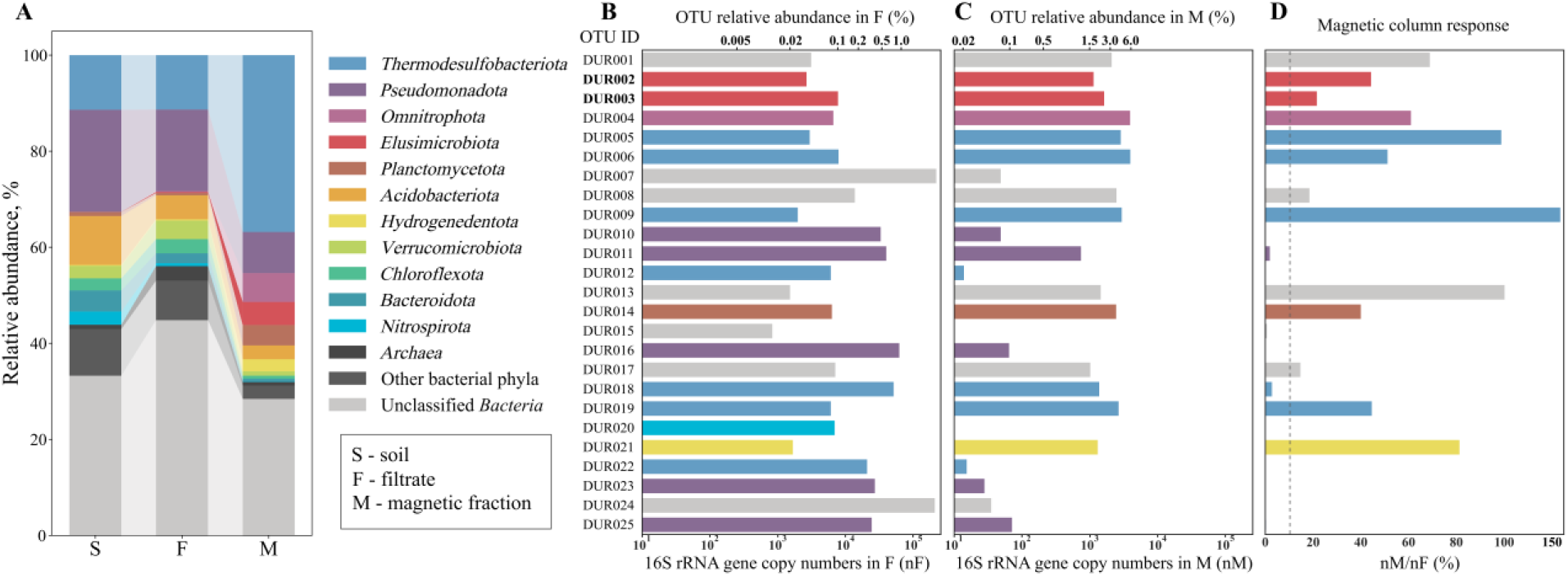
The composition of the microbial community of the analyzed black bog soil. (**A**) The microbial community composition in the initial soil sample from the microcosm (S), after filtration (F), and after enrichment with a magnetic column (M). (**B**) Relative abundance of the top 25 zOTUs in filtrate. (**C**) Relative abundance of the top 25 zOTUs in the magnetic-enriched fraction. (**D**) Magnetic column response values of the top 25 zOTUs.

The bacterial composition was similar in the F and S samples, with the following dominant groups: *Pseudomonadota* (17.0%), *Thermodesulfobacteriota* (11.3%), and *Acidobacteriota* (5.0%). The *Archaea* percentage significantly increased from 0.9% to 3.0% after the filtration step. The small size of some nano-sized archaeal cells probably allows them to pass through the filter pores more easily than the more massive bacterial cells [54]. The M and F community compositions differed considerably, as the *Archaea* percentage decreased by up to 0.8% after magnetic enrichment. The predominant phyla were *Thermodesulfobacteriota* (36.8%) and *Pseudomonadota* (8.6%). An abundance of some minor soil community phyla increased significantly after magnetic enrichment, with the relative abundance of *Omnitrophota* increasing 272 times (from 0.022% to 6.00%), *Elusimicrobiota* almost 200 times (from 0.025% to 4.814%), *Hydrogenedentota* 12.8 times (from 0.197% to 2.53%), and *Planctomycetota* more than 5 times (from 0.81% to 4.23%). This considerable enrichment of these four bacterial phyla was very likely due to MTB separation, as MTB were previously detected in all of them [15, 24, 55]. In all three samples, a large fraction of bacterial zOTUs was not classified at the phylum level. The share of unclassified zOTUs was 32% in the S community, 44% in the F community, and 28% in the M community.

Electron microscopy revealed cells containing electron-dense magnetosomes in the enriched magnetic fraction (Fig. 2). Two cell types were distinguished by morphology: vibrio cells 3.5-4.0 μm long and 0.4–0.5 μm wide (Fig. 2 A-C), and bacilli 1.1-1.6 μm long and 0.6 μm wide (Fig. 2 D-F). Vibrio synthesized elongated bullet-shaped magnetosomes up to 110 nm long (Fig. 2A) and drop-shaped magnetosomes up to 60 nm in length (Fig. 2 B, C). The rods synthesized fewer than 10 elongated tooth-shaped and bullet-shaped magnetosomes per cell measuring 50–90 nm in length. None of the detected morphotypes contained magnetosomes organized in the chains typical of MTB.

**Fig. 2:**
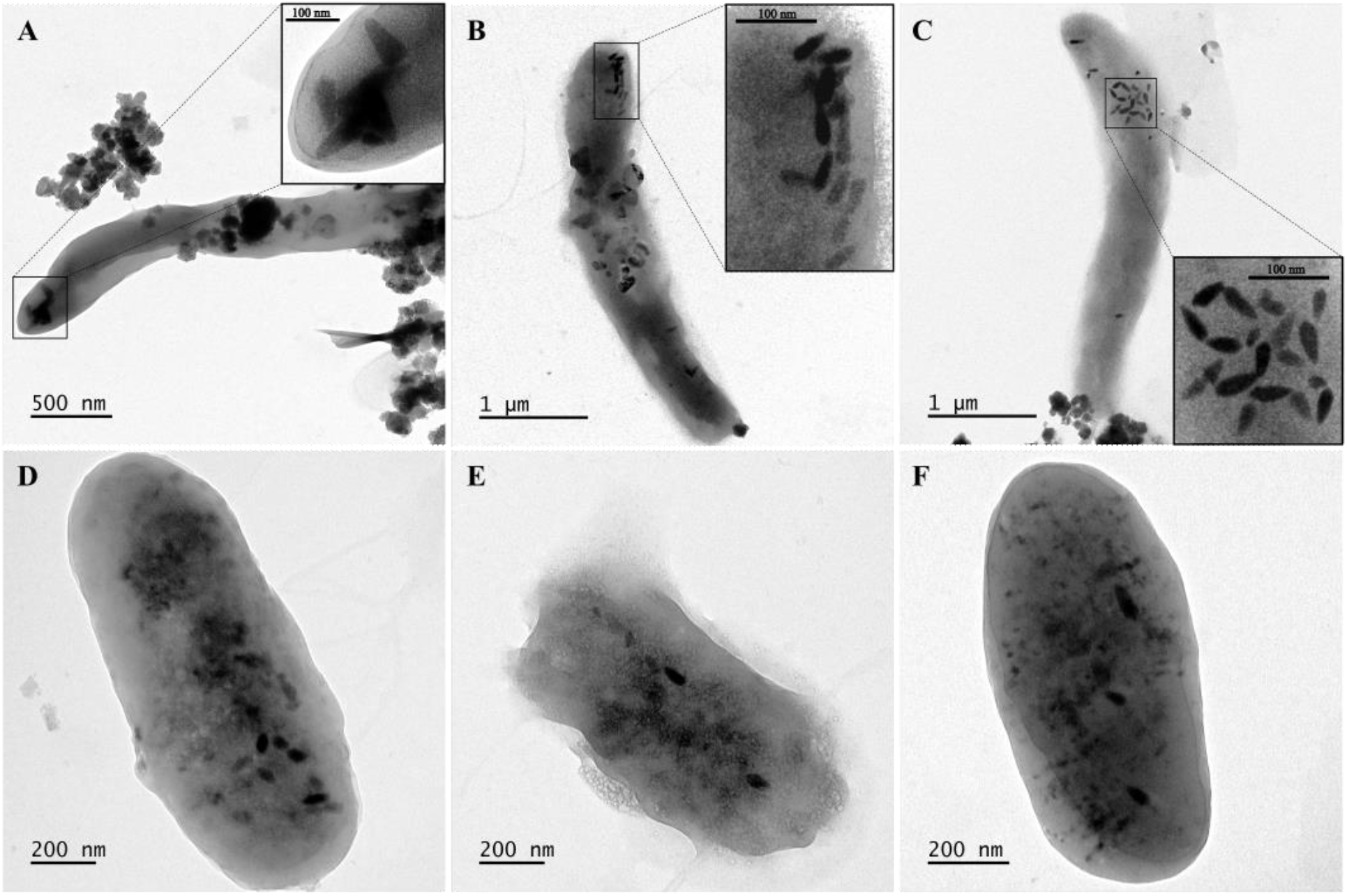
TEM images of enriched magnetic cells. (A – C) vibrio-shaped cells, (D-F) rod-shaped cells. The enlarged area in which the magnetosomes are located is highlighted in the black square.

The changes in zOTU abundance during magnetic separation were explored by calculating the number of gene copies per gram of the samples using RT-PCR. The resulting copy numbers averaged 1.0×10^8^, 7.3×10^6^, and 6.58×10^4^ gene copies g-1, respectively, for samples S, F, and M. Thus, the magnetically enriched cell fraction accounted for approximately 0.07% of the S and 0.9% of the F communities. The zOTUs enriched after magnetic separation were identified by calculating the ratio of 16S rRNA gene copies in the magnetic fraction to the number of copies in the filtrate (nM/nF) (Fig. 1BCD, Supplementary Table S3), and 25 of the most abundant zOTUs in all samples were selected for more detailed analysis. A zOTU was considered responsive to magnetic separation when the nM/nF value was greater than 10%. Overall, 12 zOTUs that did not show a magnetic response were closely related to a known species incapable of forming magnetic particles (*Nitrosospira*, *Methyloversatilis*, etc.) [56, 57], whereas13 zOTUs (DUR001, DUR002, DUR003, DUR004, DUR005, DUR006, DUR008, DUR009, DUR013, DUR014, DUR017, DUR019, and DUR021) showed a magnetic response. Among these zOTUs, DUR005 showed a high level of similarity to known non-magnetotactic magnetic particle producers. DUR005 was closely related to *Geobacter hydrogenophilus* H2 with a similarity level of 85.5%. *Geobacter* spp. are known to biomineralize magnetic nanoparticles, which may contribute to their magnetic enrichment [58]. However, this zOTU was also close to magnetotactic *Ca*. Belliniella magnetica LBB04 [59] with a similarity level of 85.4%; therefore, we cannot rule out DUR005 as a MTB. DUR006 was similar to the *Ca*. Belliniella magnetica LBB04 at 97.2% and most likely belongs to MTB. *Ca*. Belliniella magnetica LBB04 is a 2.5 μm long rod with elongated disorganized magnetosomes [27] and a morphology similar to the cells shown in Fig.2 E-F. DUR006 was the most abundant (5.9%) in the M fraction. Microscopy observation revealed many cells similar in morphology to *Ca*. Belliniella magnetica LBB04, suggesting a connection of the morphotypes from Fig.2 E-F with DUR006. Two zOTUs, DUR009 and DUR019, belonged to phylum *Thermodesulfobacteriota*, with a high level of similarity to *Ca*. Belliniella magnetica LBB04 (84.1% and 88.6%, respectively). The remaining zOTUs belonged to the phyla *Omnitrophota, Planctomycetetota*, *Hydrogenedentota*, and *Elusimicrobiota*, where the presence of MTB was reported previously. The taxonomic position of four zOTUs (DUR001, DUR008, DUR013, and DUR017) could not be reliably determined at the phylum level.

Among the most represented zOTUs in sample M were DUR002 and DUR003, which accounted for about 4.2% of all amplicons in this community. DUR002 and DUR003 had about 89% similarity to the cultivated *Endomicrobium proavitum* Rsa215 of the phylum *Elusimicrobiota*. Until recently, only one draft genome of MTB was found in this phylum [24]. The rather impressive enrichment of the DUR002 and DUR003 suggested the presence of magnetic particles in them, so they were subjected to further investigation.

### Genome reconstruction and phylogenomic analyses

Metagenomic sequencing of the enriched magnetic fraction resulted in 106 276 479 short paired-end raw sequence reads (16.1 Gb) and 299 619 ONT-passed long reads (2.1 Gb). Hybrid metagenomic assembly of long and short reads and genome reconstruction produced two metagenome-assembled genomes (MAGs) that met the GTDB criteria: completeness ≥ 50% and contamination < 5% [60]. The 16S rRNA sequences detected in these genomes corresponded to the DUR002 and DUR003 zOTUs. The DUR002 genome was reconstructed with an assembly completeness of 94.4%, with a length of 3.4 Mb (N_50_ – 185 704 bp) and GC composition of 39.8% (Supplementary Table S4). The DUR003 genome was 2.9 Mb long, with N50 of 46 489, GC composition of 52.8%, and assembly completeness of 75.8%.

According to GTDB r207, the genomes of DUR002 and DUR003 are affiliated with the *Elusimicrobiota* phylum. These genomes, together with 217 *Elusimicrobiota* genomes, were subjected to phylogenetic analyses (Fig. 3, Supplementary Table S5).

**Fig. 3:**
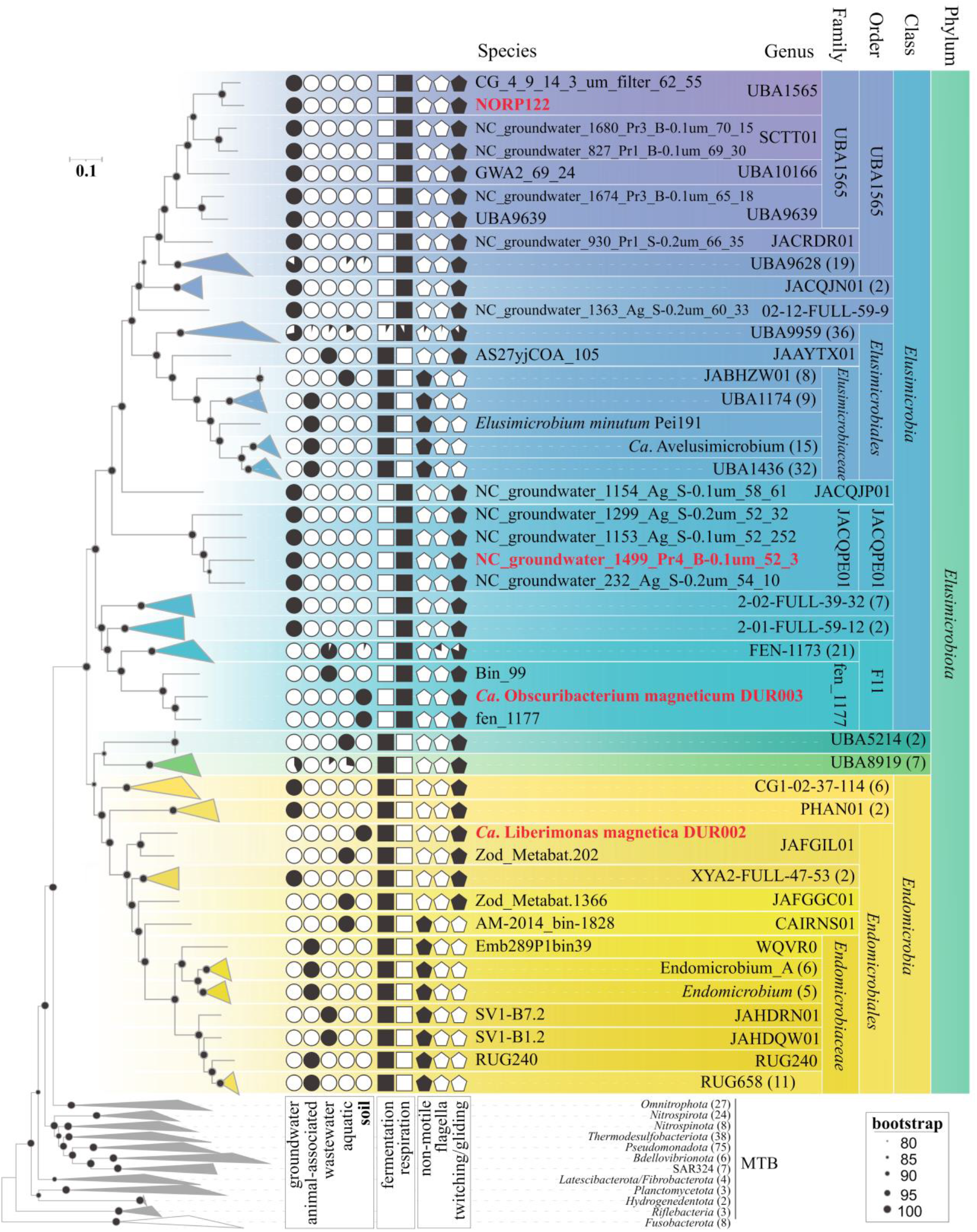
Maximum-likelihood phylogenomic tree of all known *Elusimicrobiota*, all MTB genomes previously known or obtained in this work. The tree was inferred from 120 concatenated bacterial single-copy marker proteins constructed with the evolutionary model LG + F + I + G4. Branch supports were obtained with 1000 ultrafast bootstraps. The scale bar represents amino acid substitutions per site. MTB genomes studied in this work are highlighted in red. Circles depict the habitat preferences of all studied genomes. Squares indicate the presence of fermentation-based or respiratory-based metabolism genes. Pentagons reflect the presence or absence of genes for a certain motility type.

The *Elusimicrobiota* phylum contains four classes: *Endomicrobia*, *Elusimicrobia*, UBA5214, and UBA8919. The DUR002 genome belongs to the *Endomicrobia* class, which contains 40 genomes, but DUR002 is the only genome containing magnetosome genes. As per GTDB, DUR002 and the closely related genome Zod_Metabat.202 [61] belong to the same family JAFGIL01. The AAI value between DUR002 and Zod_Metabat.202 was below 65%, which means that they belong to different genera [62]. The Zod_Metabat.202 genome was obtained from a freshwater sediment biome similar in its environmental parameters to the sampling site for DUR002.

According to GTDB, DUR003 belongs to the *Elusimicrobia* class, which includes 170 genomes. DUR003 was closely related to fen_1117 and Bin_99, and they belong to the same family fen_1177 of the order F11. The AAI values between these three genomes were below 65%; therefore, they could be assigned to three different genera [62] (Supplementary Table S6). The genome of fen_1177, like DUR003, originated from the peat metagenome. In addition to DUR003, two more MTB genomes belong to the *Elusimicrobia* class (discussed below). One of them, NORP122, identified as MTB previously [24], belongs to the order UBA1565, whose members are all groundwater inhabitants. The second MTB genome, NC_groundwater_1499_Pr4_B-0.1um_52_3 (henceforth “1499”) identified in this study, belongs to the order JACQPE01, also represented by groundwater inhabitants.

On the phylogenomic tree, all the studied MTB genomes clustered with the genomes of free-living *Elusimicrobiota*. By contrast, animal-associated genomes form separate, well-traced clades affiliated with the family *Endomicrobiaceae* (*Endomicrobia*) and with the family *Elusimicrobiaceae* (*Elusimicrobia*). No MTB representatives were detected among the animal-associated genomes or in the UBA5214 and UBA8919 classes.

The species tree and GTDB taxonomy indicated that the DUR002 and DUR003 genomes are distant from all cultivated or validly published *Elusimicrobiota* species. Due to their position on the phylogenomic tree, and according to AAI data, we assigned DUR002 and DUR003 to novel genera and novel families and proposed the following names: *Candidatus* Liberimonas magnetica and *Ca*. Obscuribacterium magneticum, respectively.

### Metabolic potential of magnetotactic *Elusimicrobiota*

Analysis of the *Elusimicrobiota* MTB genomes reveals a considerable diversity of their metabolic capacities (Supplementary Tables S7-S12). Based on the completeness of the carbon and energy pathways, HCA allowed sorting of 219 available MAGs, including DUR002 and DUR003, into two major groups: exploiters of fermentation-based and respiratory-based metabolism (Supplementary Fig. S1). Interestingly, the genomes containing magnetosome genes were found in both groups: DUR003, NORP122, and 1499 appear to have respiration capacities, whereas DUR002, to our knowledge, is the first MTB with a fermentation-based metabolism described to date (see the description below).

DUR002 lacks a complete TCA, NADH dehydrogenase (Complex I), and all but one component, (Complex V, F-type ATPase) from the oxidative phosphorylation electron transport chain, suggesting it is an obligate fermenter. Fermenting *Elusimicrobiota* almost always have Complex V; however, its role remains uncertain [19]. DUR002 contains a nearly complete glycolysis (Embden-Meyerhof pathway) and pentose-phosphate pathway and has the capacity to produce lactate and acetate as fermentation products. No enzymes suggestive of ethanol or malate fermentation were found.

The genome of DUR002 encodes numerous glycoside hydrolases and glycosyl transferases, indicating an ability to utilize various external polysaccharides. DUR002 also has the potential for autotrophic growth with hydrogen and carbon dioxide via the Wood-Ljungdahl pathway. The energetic conservation associated with carbon dioxide reduction to acetyl-CoA likely relies on the Rnf (sodium-motive ferredoxin:NAD+ oxidoreductase) complex, as predicted previously for some lineages of groundwater-associated *Elusimicrobiota* [19].

DUR002 is predicted to lack nitrite or nitrate assimilation ability, as no nitrate or nitrite transporters were found in the genome. Unlike some previously described *Elusimicrobiota*, DUR002 is also incapable of nitrogen fixation [19]. Nitrogen is most likely derived from extracellular ammonium, as ammonium transporters of Amt/Mep/Rh family are encoded in the genome. Sulfur is likely assimilated through sulfate reduction through the sulfate adenylyltransferase CysNDC.

Despite its lower genome completeness than in DUR002, some predictions can be made regarding DUR003 metabolism. Identification of several parts of the electron-transport chain, including Complex I (NADH:quinone oxidoreductase), Complex III: Cytochrome bd ubiquinol oxidase, and Complex IV High affinity: Cytochrome bd ubiquinol oxidase, suggested a respiratory metabolism. DUR003 likely primarily accepts oxygen at low concentrations (microoxic) as a terminal electron acceptor. Among alternative reductases, only nitric oxide reductase NorBC was found, indicating the presence of an incomplete denitrification pathway. The TCA cycle is incomplete (only isocitrate dehydrogenase, 2-oxoglutarate/2-oxoacid ferredoxin oxidoreductase, citrate synthase, and aconitate hydratase are present), suggesting that energy may be derived from an inorganic substrate (lithotrophy) rather than from organic compounds. However, the substrate is difficult to predict due to genome incompleteness. No key genes for either sulfur-iron or CO oxidation were found. Among hydrogenases, DUR003 contained one subunit (alpha) of a membrane-bound hydrogenase [EC:1.12.7.2], an enzyme often involved in other bacteria in H2 production rather than oxidation. Therefore, whether H2 can serve as a substrate also remains uncertain.

No nitrate or nitrite reduction and no nitrogen fixation were found in DUR003. As with DUR002, an Amt/Mep/Rh family transporter is encoded, suggesting that nitrogen is derived from ammonium. Sulfate can be imported; however, no genes for assimilatory sulfate reduction were found. Dissimilatory sulfate reduction, sulfur oxidation, and the SOX system are absent.

Magnetosome formation is linked to cellular iron transport and homeostasis. Therefore, we also assessed genes associated with iron metabolism in potentially magnetotactic *Elusimicrobiota*. Both DUR002 and DUR003 encode numerous iron transport and associated regulatory genes, mostly linked to siderophore-mediated transport. Thus, DUR003 has several putative transport systems for pyoverdine-(PvdRT) and enterobactin-like (EntS) siderophores. Its genome also has 9 genes for a putative regulator FecR that functions as a sensor for iron (III) in *E. coli*.

DUR002 contains several systems for putative pyoverdine-like siderophores (FpvE, PvuERT). Iron storage likely occurs through ferritins, as the genome contains at least three ferritin-domain-containing proteins. At least two genes encoding Fur-family transcriptional repressors are found in DUR002, suggesting their role in the regulation of iron homeostasis.

### Genome-based prediction of motility

Magnetotaxis is a combination of passive alignment to the external magnetic field and active movement along the field lines in a direction selected based on common aero- or chemotactic sensing. Hence, active motility is key to exploiting the benefits of magnetotaxis in native environments. Indeed, all previously described MTB have flagella and are highly motile [7]. The knowledge is scarce regarding motility in *Elusimicrobiota* representatives and is based on isolated gut or intracellular symbiotic species, which were described as non-motile [16, 18]. Unlike the capillary racetrack method broadly applied previously to isolate MTB, MTB-CoSe separation requires only the presence of magnetic minerals within the cells, without exploiting active movement. Therefore, the isolation method does not imply that these bacteria are able to move like other MTB and raises the question whether magnetotactic *Elusimicrobiota* are actually able to move. We answered this question by reconstructing their potential motility based on sequenced genomes.

HCA analysis based on the abundance of motility genes (KEGG BRITE: Bacterial motility proteins) revealed several major *Elusimicrobiota* groups whose clustering might correlate with the ability to move: (i) non-motile, (ii) motile by means of flagella, and (iii) supposedly exploiting twitching motility by Type IV pili (T4P) (Supplementary Fig. S2). Most symbiotic species are likely non-motile, as they lack flagellum structural genes and related regulatory proteins as well as two-component signal transduction systems regulating chemotaxis (Che) and methyl-accepting chemotaxis proteins (MCP). Notably, they possess numerous homologues of the type IV pili apparatus assembly proteins PilABCDQME, with the pilin protein PilA being the most abundant. However, their lack of regulatory components for the twitching motility apparatus (PilGHR) and for chemotaxis protein genes suggests a different role for Type IV pili in these species, apart from motility (e.g., DNA uptake or protein secretion) [63]. Only a few *Elusimicrobiota* genomes contain the complete set of genes for building a flagellum apparatus, but most free-living representatives, including the supposedly magnetotactic species, appear to have genes involved in chemotaxis, methyl-accepting receptors, and Type IV pili assembly, suggesting potential twitching motility in response to stimuli. Thus, the genome of DUR002 encodes homologues of CheAWCDRBY, MCP, and the sensory regulator, ChpA. The genome of DUR003 encodes homologues of CheAWDBY, but no MCP are found, presumably due to the lower genome completeness. Both contain the homologues of the T4P apparatus proteins PilTABCDQM. Interestingly, despite the absence of structural flagellum proteins, the magnetotactic *Elusimicrobiota*, as with most of the free-living species, contain numerous homologues of the genes for *motAB* (motor switch in the flagellum stator), *flbD* (transcriptional activator of the flagellum genes), and *fliG* (motor switch in the flagellum rotor). The DUR003 genome also encodes a gliding motility-associated transport system, GldAFG. Both MotAB-like and FliG proteins provide a protonmotive force for driving gliding motility in some bacteria [64]. Therefore, at least in DUR003, MotAB might be functionally linked to a Gld system. However, in *Elusimicrobiota* lacking the Gld system, including DUR002, the role of MotAB is unclear. Previously, transposon-mediated mutation of these genes was shown to affect twitching motility in *Pseudomonas aeruginosa*, so it might also be functionally involved in twitching [65]. However, the exact role in twitching in *Elusimicrobiota* remains enigmatic.

Further investigation is needed to establish whether twitching or gliding motility occurs in the magnetic *Elusimicrobiota*. Nonetheless, our predictions suggest that magnetotaxis might be linked to motility types other than flagella-based.

### Magnetosome gene cluster reconstruction

The DUR002 and DUR003 genomes reconstructed in this work revealed putative magnetosome gene clusters (MGCs) (Fig. 4A). We also used the MagCluster tool to search for magnetosome synthesis genes in the entire *Elusimicrobiota* set used in this study. This allowed discovery of a MGC in one more genome, named 1499 (mentioned above). The NORP122 genome also revealed a second contig with MGC, in addition to the existing one.

**Fig. 4:**
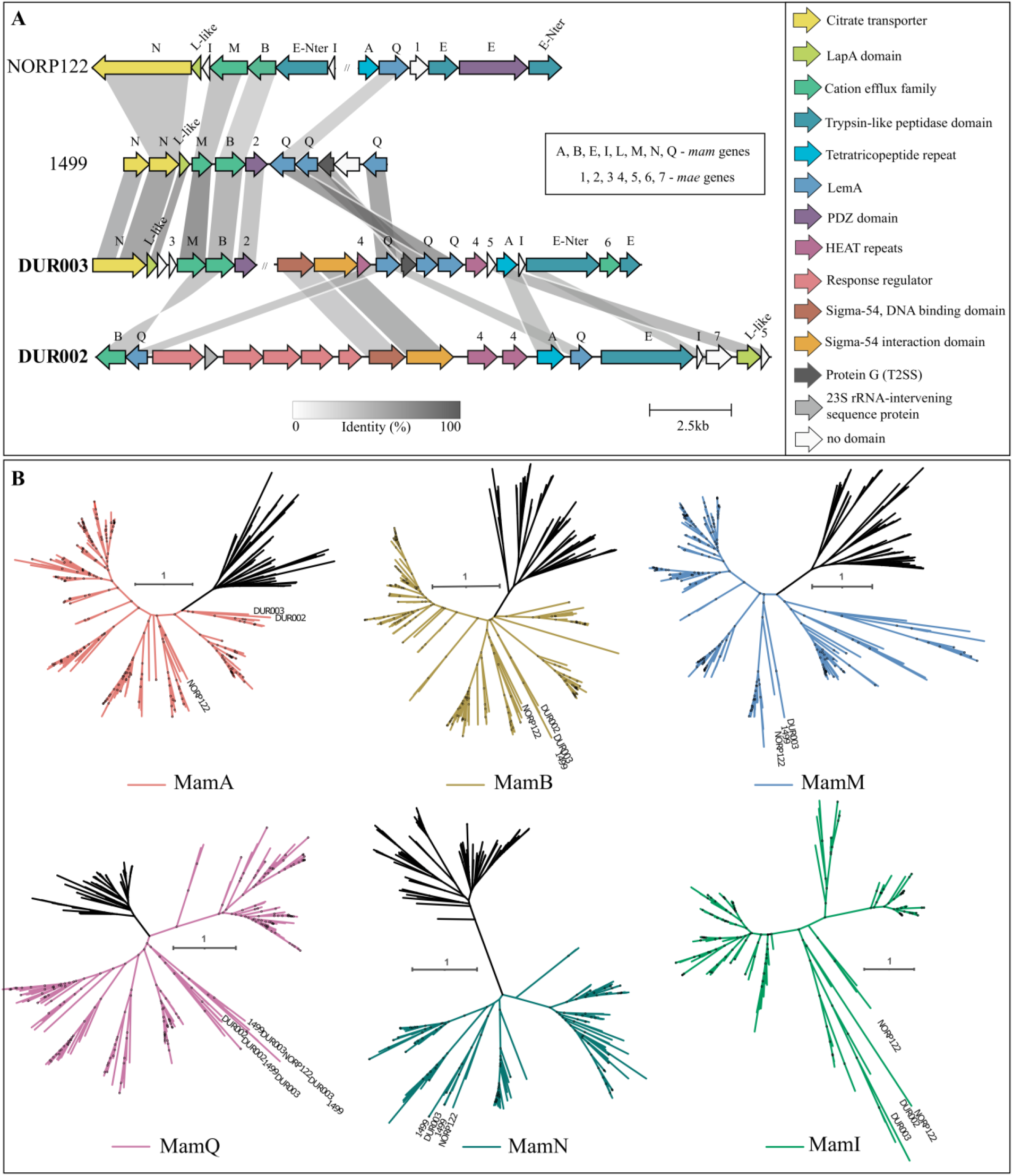
Magnetosome gene clusters of the investigated *Elusimicrobiota* genomes. (**A)** Comparison of the MGC regions in the *Elusimicrobiota* MTB genomes. The genes are colored according to the conserved domains they include. (**B**) Maximum-likelihood phylogenetic trees based on Mam protein sequences of studied and other known MTB along with their non-MTB homologues. Trees were reconstructed with LG + F + I + G4 substitution model. Branch supports were obtained with 1000 ultrafast bootstraps. The scale bar represents amino acid substitutions per site. Full trees can be found in figshare data [66].

The four studied MGCs contained putative *mam* genes, including *mamA*, -*B*, -*M*, -*I*, -*Q*, -*E*, -*L-like*, and *-N*. The protein sequences of these genes from *Elusimicrobiota* and other known MTB, along with their homologues from non-MTB, were used to construct phylogenetic trees (Fig. 4B, figshare supplementary data [66]). The Mam protein sequences of the *Elusimicrobiota* MGCs almost always clustered with each other and with those of *Fibrobacterota* and *Riflebacteria*. The exception was NORP122, whose MGC sequences, in some cases, clustered with other MTB representatives. We also noted 3 copies of the MamQ protein in the MGCs of the 1499 and DUR003.

The conserved domains of *Elusimicrobiota* MGCs were determined using the PFAM and COG databases (Supplementary Table S13). *Elusimicrobiota* MGC sequences have been found to contain the same domains as the sequences of the orthologous Mam proteins in model MTB *Magnetospirillum gryphiswaldense* MSR-1, [9]. Notably, the magnetochrome domains required to maintain iron redox balance in magnetosomes [67] were detected only in NORP122 MamE and MamN proteins; 1499, DUR002 and DUR003 proteins did not contain these domains. In addition, none of the investigated MGCs contained sequences of the filamentous actin-like protein MamK responsible for alignment of magnetosomes into chains [68]. Apart from *mam* genes, the MGCs of *Elusimicrobiota* contained no other genes with known involvement in magnetosome synthesis (*mad*, *man*, *mms*, etc.).

The *Elusimicrobiota* MGCs contained several genes specific to this group. We propose the name *mae* for these genes (**ma**gnetosome ***E**lusimicrobiota*), as they have either not been found before or were found in a few MTB but have not been named (Supplementary table S13). The *mae1, −3*, and *-7* genes are unique. They do not have detected domains with PFAM and COG in their protein sequences, nor do they share any similarities (e-value cutoff 1e^-05^) with any other genes in the NCBI database. The *mae2* and *mae6* genes contained the PDZ domain and cation efflux family domain, respectively, according to PFAM. However, in phylogenetic trees, these protein sequences did not cluster with the known MamE, MamB, or MamM sequences having the same domains [9]. Both *mae2* and *mae6* are likely genes specific to *Elusimicrobiota* that perform functions similar to those performed by known Mam proteins with the same domains. The *mae4* and *mae5* genes share no similarities (e-value cutoff 1e^-05^) with non-MTB genes in the NCBI database. They are similar to *Bdellovibrionota* and *Planctomycetota* (45% of identity) MTB genes and have never been named before.

The MGCs of 1499, DUR002 and DUR003 contain magnetosome genes interspersed with genes similar to those involved in other processes occurring in both MTB and non-MTB. Thus, in 1499 and DUR003, the magnetosome genes neighbor with the type II secretion system protein GspG. DUR002 and DUR003 have genes with Sigma-54, DNA binding, and interaction domains, and DUR002 also has several response regulator genes in its MGC.

## DISCUSSION

Several methods have already been proposed for soil microcosm enrichment by subjecting them to various physical and chemical stresses [69–71]. Unlike these other methods, magnetic enrichment does not require any change in the environment beforehand and promotes the selection of rare taxa representatives with different metabolic functions. Therefore, like magnets that can remove a needle from a haystack, magnetotactic bacteria can help study the rare biosphere.

Studying the rare biosphere is challenging and even more complex in soils, as soils have enormous microbial diversity [72]. Magnetotactic bacteria are minor components of microbial communities, and some belong phylogenetically to poorly investigated phyla [15, 24]. Studies on the presence of MTB in soils are scarce, and only recently have several MTB genomes been detected in acidic peatlands [15], suggesting a potentially widespread MTB occurrence in different types of waterlogged soils, in addition to water habitats [15].

In this work, we carried out 16S rRNA sequencing of peaty black bog soil (S), its filtrate (F), and magnetically enriched bacterial community (M). The abundance of some representatives from poorly studied phyla, such as *Hydrogenedentota*, *Elusimicrobiota*, and *Omnitrophota*, increased in the M compared to the S communities. In the M community, 28% of reads were not classified at the phylum level, suggesting that these are new representatives of the rare biosphere. In addition, among the top 25 zOTUs, 13 had a magnetic response and accounted for about 45% of M, but less than 0.1% of S. Thus, magnetic enrichment using the MTB-CoSe method allowed high-coverage sequencing and study of genomes containing these zOTUs.

*Elusimicrobiota* is a poorly studied phylum with animal-associated and free-living representatives. Previously, magnetosome gene clusters were discovered only in a single genome, NORP122, belonging to *Elusimicrobiota*. However, that genome contained only a short MGC fragment comprising 7 magnetosome synthesis genes, raising doubts about considering NORP122 as an MTB [24]. In this work, we reconstructed two MTB genomes from *Elusimicrobiota* obtained from black bog soil, and we found one more MTB genome affiliated with this phylum in the NCBI database. All these results indicated that the presence of MGCs in the *Elusimicrobiota* was not an assembly error.

Analysis of MGCs of all four *Elusimicrobiota* MTB showed a rather low identity (usually between 25-35%) between each MGC protein sequence. The possible reason may be that the last common ancestor of all *Elusimicrobiota* had MGC, and its inheritance occurred vertically with multiple losses. This assumption is also supported by the fact that MTB on the *Elusimicrobiota* species tree do not form a separate clade but are spread throughout the tree (Fig. 3). However, another possibility is that some of *Elusimicrobiota* MGCs were acquired by horizontal gene transfer from MTB of other phyla. Thus, more data are required to clarify how MGCs evolved in this phylum. A detailed analysis of MGCs identified a second contig with magnetosome synthesis genes in NORP122. We revealed several *mam* genes, many of which are essential for magnetosome synthesis in other MTB: *mamA*, -*B*, -*M*, -*I*, -*Q*, -*E*, -*L-like*. However, no determinants of the magnetosome chain assembly *(mamK* or *mad28)* [68, 73] were found in these genomes, suggesting that magnetosomes in *Elusimicrobiota* MTB most likely do not form chains.

We also found several genes unique to MTB of this phylum and named them *mae* genes. Further investigations are needed to determine precisely how these genes contribute to magnetosome synthesis, size, and shape. Moreover, we only found magnetochrome domains, which are responsible for the iron redox balance statement in magnetosomes [67], in one of four *Elusimicrobiota* MGCs, NORP122. The absence of these domains in the other three MGCs suggests that these MTB can use a different mechanism to control the iron redox state during magnetosome synthesis. Another possible explanation is that these magnetosomes are composed of different mineral than magnetite or greigite. Further studies on magnetosome morphology in *Elusimicrobiota* MTB will help shed light on this issue.

Metabolic analysis revealed that DUR002 is, to our knowledge, the first obligate fermenter among MTB. This type of metabolism probably provides advantages in the anaerobic conditions of waterlogged bog soils. Genes for twitching and gliding motility, but not for flagella formation, were also detected in the new genomes, making them exceptional among MTB, as all other known representatives are motile by means of flagella. We speculate that MTB with twitching and gliding motilities were not found before because common techniques used for MTB separation were based on active fast movement enabled by flagella. In this work, we used a recently developed MTB-CoSe separation technique that does not depend on active cell movement and requires only the presence of magnetic particles within the cells. Most previous MTB studies were also conducted in aquatic habitats where flagellum-driven motility dominates. As soils have solid surfaces, surface-associated motility types, such as twitching and gliding, prevail there [74]. Therefore, magnetotaxis linked to surface-associated motility might be beneficial in soil environments.

We also detected response regulator genes consisting of CheY-like receiver domains in the DUR002 MGC. Previous reports have indicated that CheY helps switch the rotation direction of the flagellar motor [75]. We assume that the colocalization of these genes with magnetosome synthesis genes is not accidental, and that all these genes could be involved in magnetochemotaxis. However, magnetosomes may possibly perform other functions, such as elimination of toxic reactive oxygen or iron sequestration and storage, where they act as electrochemical batteries [15].

Further study of magnetotactic bacteria from the rare biosphere, including bacteria from the phylum *Elusimicrobiota*, is necessary to understand magnetosome formation and function. The findings reported here raise questions about the mechanisms of magnetotaxis without flagella and actin-like MamK protein, the process of magnetosome biomineralization in the absence of magnetochrome proteins and magnetosome functions in obligately anaerobic bacteria.

### Candidatus description

#### *Candidatus* Obscuribacterium

Obscuribacterium (Ob.scur.i.bac.te’ri.um. L. masc. adj. *obscurus*, dark; N.L. neut. n. *bacterium*, a rod; N.L. neut. n. *Obscuribacterium*, a bacterium found in the dark)

#### *Candidatus* Obscuribacterium magneticum

Obscuribacterium magneticum (mag.ne’ti.cum. L. neut. adj. *magneticum*, magnetic)

Potentially has a respiratory-based metabolism. No nitrate or nitrite reduction; no nitrogen fixation. Sulfate can be imported, but no genes for assimilatory sulfate reduction were found. Dissimilatory sulfate reduction, sulfur oxidation, and the SOX system are absent. Supposedly capable of twitching motility. Collected on a magnetic column from waterlogged soil of the Durykino ravine. The reference strain is DUR003. The genome reference sequence of DUR003 is <u>JAJAPZ000000000</u>. G+C content 52.84%

#### *Candidatus* Liberimonas

Liberimonas (Li.be.ri.mo’nas. L. masc. adj. *liber*, free; L. fem. n. *monas*, unit, monad; N.L. fem. n. Liberimonas, a free monad)

#### *Candidatus* Liberimonas magnetica

Liberimonas magnetica (mag.ne’ti.ca. L. fem. adj. *magnetica*, magnetic)

Potentially has a fermentation-based metabolism. Has the capacity to produce lactate and acetate as fermentation products. Has the potential for autotrophic growth with hydrogen and carbon dioxide via the Wood-Ljungdahl pathway. Predicted unable to assimilate nitrite or nitrate and unable to fix nitrogen. Sulfur is likely assimilated through sulfate reduction. Supposedly capable of twitching motility. Was collected on magnetic column from waterlogged soil of the Durykino ravine. The reference strain is DUR002. The genome reference sequence of DUR002 is <u>JAJAPY000000000</u>. G+C content 39.76%

## Data availability

Raw reads of metagenomic 16S rRNA sequencing of soil, filtrate and magnetic fraction were deposited in the NCBI Sequence Read Archive under the accession numbers SRR19138506, SRR19138505 and <u>SRR19078916</u>. DUR002 and DUR003 genome sequences have been deposited in GenBank under the accession numbers JAJAPY000000000 and JAJAPZ000000000 (BioProject number PRJNA769424). The raw metagenomic read data have been deposited in the NCBI Sequence Read Archive under the accession numbers <u>SRR16235686</u> and <u>SRR16235685</u>. All data generated and analyzed in this study are also available in figshare [66] and in the supplementary information accompanying this paper.

## ACKNOWLEDGMENTS

We thank Prof. Aharon Oren for his expert guidance in nomenclature. We also thank Prof. Jillian Banfield and Banfield Lab for the providing additional information about 1499 genome. Bioinformatic analyses were performed using computing resources at the Core Research Facility ‘Bioengineering’ (Research Center of Biotechnology RAS) and SciBear OU (https://sci-bear.com/). TEM studies were carried out at Electron microscopy laboratory of Moscow State University Biology Faculty. The reported study was funded by Russian Foundation for Basic Research, project number 20-34-90116.

