## Supplementary Table S1 for "Looking for a needle in a haystack: magnetotactic bacteria help in “rare biosphere” investigations": Elusi_supp_final.pdf

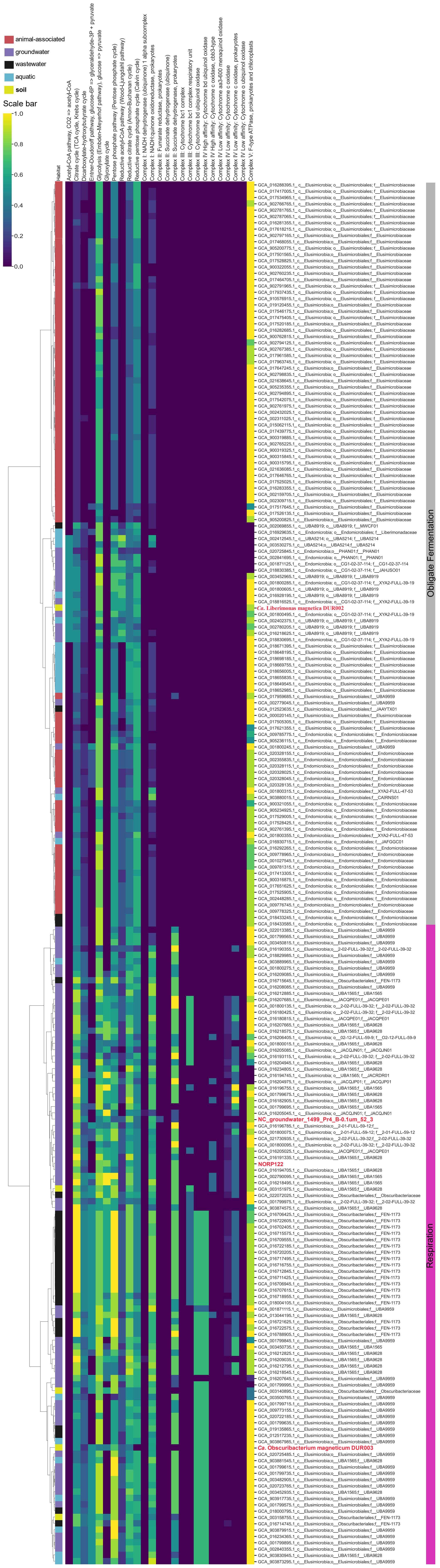

**Supplementary Figure S1.** Hierarchical clustering analysis (HCA) of Elusimicrobia genomes based on the completeness (estimated using DRAM) carbon and energy metabolism pathways. Scale bar indicates the pathway completeness. The MTB genomes are highlighted in red.

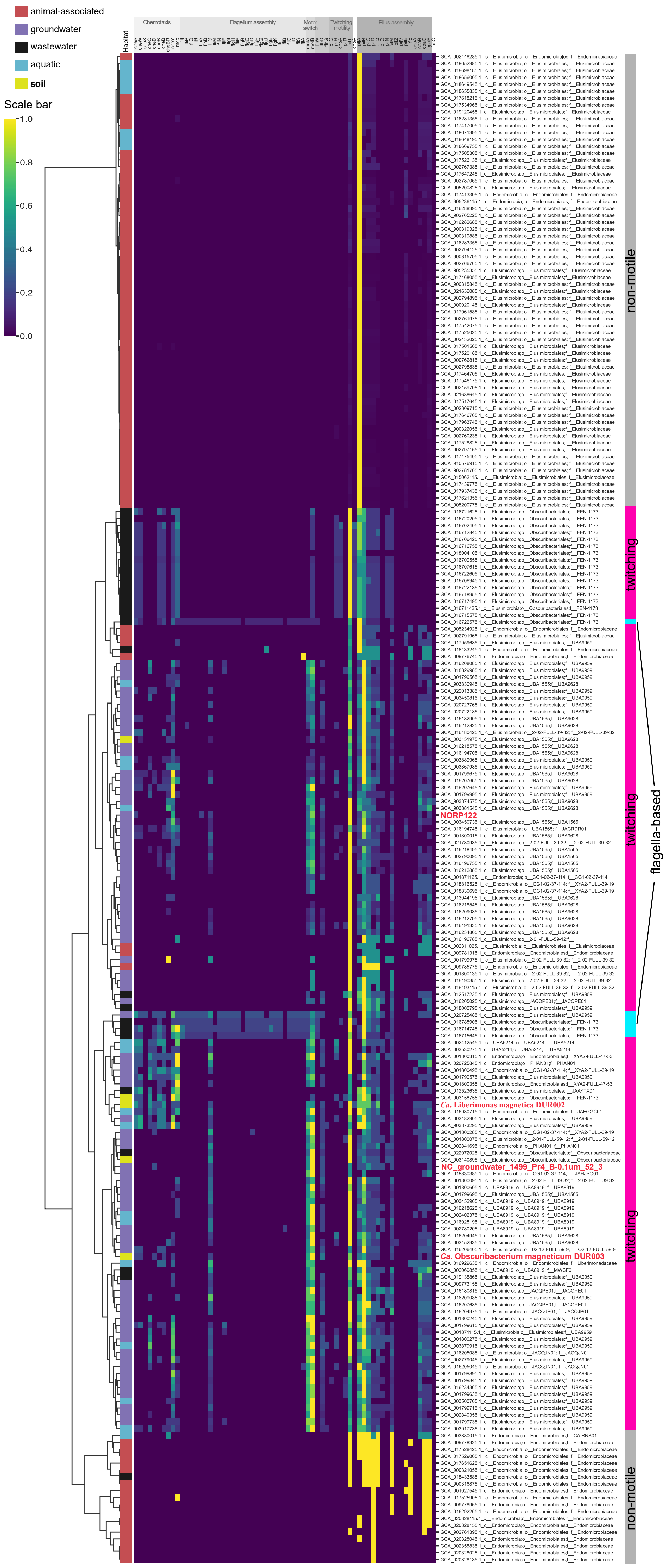

**Supplementary Figure S2.** Hierarchical clustering analysis (HCA) of Elusimicrobia genomes based on the abundance of chemotaxis and motility-related genes. Scale bar indicates the standardized number of genes. The MTB genomes are highlighted in red.

**Supplementary Table S1.** Results of the physical and chemical parameters of the analyzed microcosm.

| № | Measurement, unit of measurement | Result | № | Measurement, unit of measurement | Result |
|---|---|---|---|---|---|
| **1** | Conductivity, uS/cm | 35.0 ± 3.4 | **20** | Li, mg/dm$^3$ | < 0.01 |
| **2** | pH | 8.5 | **21** | Mg, mg/dm$^3$ | 10.7 ± 1.6 |
| **3** | $NO_3^-$, mg/dm$^3$ | < 0.1 | **22** | Mn, mg/dm$^3$ | 0.021 ± 0.006 |
| **4** | $NO_2^-$, mg/dm$^3$ | < 0.1 | **23** | Cu, mg/dm$^3$ | 0.0036 ± 0.0014 |
| **5** | $SO_4^{2-}$, mg/dm$^3$ | 20.7 ± 2.7 | **24** | Mo, mg/dm$^3$ | < 0.001 |
| **6** | $PO_4^{2-}$, mg/dm$^3$ | < 0.1 | **25** | As, mg/dm$^3$ | < 0.005 |
| **7** | $F^-$, mg/dm$^3$ | 0.17 ± 0.02 | **26** | Na, mg/dm$^3$ | 3.9 ± 0.6 |
| **8** | $Cl^-$, mg/dm$^3$ | 5.75 ± 0.75 | **27** | Ni, mg/dm$^3$ | < 0.001 |
| **9** | Al, mg/dm$^3$ | 0.154 ± 0.039 | **28** | Pb, mg/dm$^3$ | < 0.003 |
| **10** | Ba, mg/dm$^3$ | 0.052 ± 0.010 | **29** | Se, mg/dm$^3$ | < 0.005 |
| **11** | Be, mg/dm$^3$ | < 0.00010 | **30** | S, mg/dm$^3$ | 6 ± 1.1 |
| **12** | B, mg/dm$^3$ | < 0.01 | **31** | Ag, mg/dm$^3$ | < 0.005 |
| **13** | V, mg/dm$^3$ | 0.0046 ± 0.0012 | **32** | Sr, mg/dm$^3$ | 0.169 ± 0.034 |
| **14** | Fe, mg/dm$^3$ | 0.5 ± 0.08 | **33** | Sb, mg/dm$^3$ | < 0.005 |
| **15** | Cd, mg/dm$^3$ | < 0.0001 | **34** | Ti, mg/dm$^3$ | 0.0060 ± 0.0024 |
| **16** | K, mg/dm$^3$ | 1.6 ± 0.24 | **35** | P, mg/dm$^3$ | 0.2 ± 0.07 |
| **17** | Ca, mg/dm$^3$ | 82 ± 12 | **36** | Cr, mg/dm$^3$ | < 0.001 |
| **18** | Co, mg/dm$^3$ | < 0.001 | **37** | Zn, mg/dm$^3$ | < 0.005 |
| **19** | Si, mg/dm$^3$ | 14.5 ± 2.2 | **38** | $H_2S$, mg/dm$^3$ | 115.2 |

**Supplementary Table S2.** Results of 16S rRNA sequencing and alpha-diversity statistics for soil microcosm, filtrate, and magnetic enrichment fraction.

| Sample | reads | zOTUs | chao1 | equitability | jost | simpson | shannon |
|---|---|---|---|---|---|---|---|
| **S (rep_1)** | 45276 | 3342 | 3342.2 | 0.846 | 525.3 | 0.0030 | 9.91 |
| **S (rep_2)** | 43474 | 3205 | 3205.3 | 0.852 | 540.2 | 0.0029 | 9.92 |
| **S (rep_3)** | 46900 | 3399 | 3399.3 | 0.852 | 557.2 | 0.0029 | 10.00 |
| **F** | 46017 | 2723 | 2723.2 | 0.834 | 380.7 | 0.0043 | 9.51 |
| **M** | 52399 | 1273 | 1273.7 | 0.668 | 63.1 | 0.0217 | 6.89 |

**Supplementary Table S3.** Statistics for the top 25 zOTUs and results of their NCBI BLAST analyses with MTB and non-MTB

| **OTU ID** | **OTU relative abundance (%)** | | | **16s rRNA gene copy numbers** | | | **nM/nF (%)** | **Magnetic column response** | **Top-hit 16S rRNA sequence, including uncultured/environmental sample (NCBI acc. #), similarity** | **Top-hit 16S rRNA sequence, excluding uncultured/environmental sample (NCBI acc. #), similarity** | **Top-hit 16S rRNA sequence of validly described bacteria (NCBI acc. #), similarity** | **Top-hit 16S rRNA sequence of MTB (NCBI acc. #), similarity** | **Phylum** |
|---|---|---|---|---|---|---|---|---|---|---|---|---|---|
| | Soil (S) | Filtrate (F) | Magnetic fraction (M) | Soil(S) | Filtrate (F) | Magnetic fraction (M) | | | | | | | |
| DUR001 | 0.001 | 0.041 | 3.158 | 1.01E+03 | 3.09E+03 | 2.09E+03 | 68.9 | TRUE | Uncultured bacterium BB-B20 (GQ844331), 84.2% | *Syntrophorhabdus* sp. TB (AB611035), 84.2% | *Vicinamibacter silvestris* Ac_5_C6 (KP761690), 80.0% | Nitrospinae bacterium nWMHbin6 (JADGCD010000264), 80.6% | Unclassified Bacteria |
| **DUR002** | 0.002 | 0.035 | 1.709 | 2.02E+03 | 2.64E+03 | 1.13E+03 | 44.3 | TRUE | Uncultured bacterium ELA_111314_OTU_1260 (KY516811), 97.5% | Elusimicrobia bacterium RIFOXYB12_FULL_50_12 (MGVK01000024), 91.9% | *Endomicrobium proavitum* Rsa215 (CP009498), 88.7% | Elusimicrobia bacterium NORP122 (NVTF01000082), 79.2% | ***Elusimicrobiota*** |
| **DUR003** | 0.009 | 0.102 | 2.446 | 9.08E+03 | 7.68E+03 | 1.62E+03 | 21.6 | TRUE | Uncultured bacterium TE1b515h12_7891 (JQ369221), 92.6% | Elusimicrobia bacterium Bin_99 (JAJVIP010000006), 88.1% | *Endomicrobium proavitum* Rsa215 (CP009498), 85.1% | Elusimicrobia bacterium NORP122 (NVTF01000082), 80.0% | ***Elusimicrobiota*** |
| DUR004 | 0.016 | 0.087 | 5.880 | 1.61E+04 | 6.55E+03 | 3.90E+03 | 61.0 | TRUE | Uncultured bacterium HDB_SIST624 (HM187352), 96.5% | *Ca.* Velamenicoccus archaeovorus LiM (CP019384), 86.1% | *Desulforegula conservatrix* Mb1Pa (AUEY01000125), 81.1% | *Ca.* Omnitrophica bacterium nDJH13bin20 (JADFXH010000013), 83.9% | *Ca.* Omnitrophota |
| DUR005 | 0.012 | 0.039 | 4.288 | 1.21E+04 | 2.94E+03 | 2.84E+03 | 98.8 | TRUE | Uncultured bacterium OTU2881 (MF452574), 98.2% | *Geobacter* sp. SVR (AP024469), 89.1% | *Geobacter hydrogenophilus* H2 (U28173), 88.5% | *Ca.* Belliniella magnetica LBB04 (MK632188), 85.4% | *Thermodesulfobacteriota* |
| DUR006 | 0.032 | 0.104 | 5.926 | 3.23E+04 | 7.83E+03 | 3.93E+03 | 51.2 | TRUE | Uncultured proteobacterium TRF-215 (JX859941), 99.4% | *Syntrophus aciditrophicus* SB (CP000252), 93.8% | *Syntrophus aciditrophicus* SB (CP000252), 93.8% | *Ca.* Belliniella magnetica LBB04 (MK632188), 97.2% | *Thermodesulfobacteriota* |
| DUR007 | 0.154 | 2.858 | 0.073 | 1.55E+05 | 2.15E+05 | 4.84E+01 | 0.0 | FALSE | Uncultured bacterium CSBC1F02 (GU127054), 90.9% | *Ignavibacterium album* JCM 16511(CP003418), 85.5% | *Ignavibacterium album* JCM 16511(CP003418), 85.5% | *Ca.* Magnetomorum sp. HK-1 (JPDT01000165), 79.3% | Unclassified Bacteria |
| DUR008 | 0.042 | 0.180 | 3.711 | 4.24E+04 | 1.36E+04 | 2.46E+03 | 18.5 | TRUE | Uncultured organism SBZO_2108 (JN530671), 94.0% | *Ca.* Sumerlaea chitinivorans BY40 (CP030759), 81.7% | *Halothermothrix orenii* H 168 (NR_074915), 81.1% | Nitrospinae bacterium nWMHbin6 (JADGCD010000264), 79.2% | Unclassified Bacteria |
| DUR009 | 0.002 | 0.026 | 4.441 | 2.02E+03 | 1.96E+03 | 2.94E+03 | 153.5 | TRUE | Uncultured bacterium pinkB.2010_8-clones-1 (KF513106), 91.9% | *Desulfatiglans parachlorophenolica* DS (AB763347), 87.5% | *Desulfatiglans parachlorophenolica* DS (AB763347), 87.5% | *Ca.* Belliniella magnetica LBB04 (MK632188), 84.1% | *Thermodesulfobacteriota* |
| DUR010 | 1.737 | 0.435 | 0.073 | 1.75E+06 | 3.27E+04 | 4.84E+01 | 0.2 | FALSE | Uncultured bacterium AN040 (GQ860021), 98.7% | Bacterium SCGC AAA018-P21 (HQ290515), 95.3% | *Azoarcus olearius* DQS-4 (EF158388), 92.5% | *Desulfamplus* sp. nJC1bin9 (JADFZQ010000075), 84.3% | *Pseudomonadota* |
| DUR011 | 0.508 | 0.524 | 1.111 | 5.13E+05 | 3.95E+04 | 7.37E+02 | 1.9 | FALSE | *Methyloversatilis* sp. LF (MK795691), 99.6% | *Methyloversatilis* sp. LF (MK795691), 99.6% | *Methyloversatilis discipulorum* FAM1 (AZUP01000001), 99.4% | *Desulfamplus* sp. nJC1bin9 (JADFZQ010000075), 85.1% | *Pseudomonadota* |
| DUR012 | 0.988 | 0.080 | 0.021 | 9.97E+05 | 6.02E+03 | 1.39E+01 | 0.2 | FALSE | Uncultured bacterium A6B8 (LN715716), 99.1% | *Ca.* Electronema nielsenii Freshwater_Gib-F5 (KP728465), 87.9% | *Simulacricoccus ruber* MCy10636 (MH094235), 87.3% | Deltaproteobacteria bacterium YD0425bin50 (PDZT01000031), 83.2% | *Thermodesulfobacteriota* |

| OTU ID | OTU relative abundance (%) | | | 16s rRNA gene copy numbers | | | nM/nF (%) | Magnetic column response | Top-hit 16S rRNA sequence, including uncultured/environmental sample (NCBI acc. #), similarity | Top-hit 16S rRNA sequence, excluding uncultured/environmental sample (NCBI acc. #), similarity | Top-hit 16S rRNA sequence of validly described bacteria (NCBI acc. #), similarity | Top-hit 16S rRNA sequence of MTB (NCBI acc. #), similarity | Phylum |
|---|---|---|---|---|---|---|---|---|---|---|---|---|---|
| | Soil (S) | Filtrate (F) | Magnetic fraction (M) | Soil(S) | Filtrate (F) | Magnetic fraction (M) | | | | | | | |
| DUR013 | 0.000 | 0.020 | 2.174 | 0.00E+00 | 1.51E+03 | 1.44E+03 | 100.2 | TRUE | Uncultured bacterium ELA_111314_OTU_6783 (KY521808), 100.0% | *Calorithrix insularis* KR (KX225427), 82.0% | *Calorithrix insularis* KR (KX225427), 82.0% | *Ca*. Belliniella magnetica LBB04 (MK632188), 79.8% | Unclassified Bacteria |
| DUR014 | 0.007 | 0.083 | 3.664 | 7.06E+03 | 6.25E+03 | 2.43E+03 | 40.0 | TRUE | Uncultured bacterium SS_LKC22_UB80 (AM490670), 98.1% | Planctomycetaceae bacterium ECT2AJA-110-A (CP030884), 83.7% | *Brevitalea deliciosa* Ac_16_C4 (NR_151988), 78.5% | Planctomycetes bacterium isolate MAG_18080_pl_157 (DUZQ01000079), 73.7% | *Planctomycetota* |
| DUR015 | 0.678 | 0.011 | 0.008 | 6.84E+05 | 8.28E+02 | 5.30E+00 | 0.6 | FALSE | Uncultured bacterium HDB_SIPC470 (HM186890), 98.5% | *Syntrophorhabdus aromaticivorans* UI (KI867150), 82.2% | *Syntrophorhabdus aromaticivorans* UI (KI867150), 82.2% | Nitrospinae bacterium nWMHbin6 (JADGCD010000264), 81.9% | Unclassified Bacteria |
| DUR016 | 2.075 | 0.815 | 0.097 | 2.09E+06 | 6.14E+04 | 6.43E+01 | 0.1 | FALSE | Uncultured bacterium JRB4-Oct14 (MH934558), 99.1% | Proteobacteria bacterium IMCC26096 (MW884648), 94.9% | *Nitrosospira tenuis* Nv1 (FOBH01000006), 91.6% | *Desulfamplus* sp. nJC1bin9 (JADFZQ010000075), 87.2% | *Pseudomonadota* |
| DUR017 | 0.005 | 0.093 | 1.525 | 5.05E+03 | 7.00E+03 | 1.01E+03 | 14.7 | TRUE | Uncultured bacterium 1338 (KY691750), 97.6% | Bacterium YC-ZSS-LKJ31 (KP174519), 81.4% | *Halochromatium roseum* JA134 (AM283535), 80.7% | Nitrospirae bacterium MYbin3 (PEAD01000007), 79.6% | Unclassified Bacteria |
| DUR018 | 0.115 | 0.674 | 2.060 | 1.16E+05 | 5.07E+04 | 1.37E+03 | 2.8 | FALSE | Uncultured bacterium GrasBac037 (KC161712), 98.9% | Syntrophaceae bacterium JGI 0000059-J07 (KJ638710), 93.8% | *Smithella propionica* R4b16 (AF482441), 92.5% | *Ca*. Belliniella magnetica LBB04 (MK632188), 93.9% | *Thermodesulfobacteriota* |
| DUR019 | 0.094 | 0.080 | 3.982 | 9.49E+04 | 6.02E+03 | 2.64E+03 | 44.6 | TRUE | Uncultured bacterium BSN022 (AB364726), 98.9% | Syntrophaceae bacterium JGI 0000059-J07 (KJ638710), 90.0% | *Syntrophus aciditrophicus* SB (CP000252), 89.3% | *Ca*. Belliniella magnetica LBB04 (MK632188), 88.6% | *Thermodesulfobacteriota* |
| DUR020 | 0.535 | 0.091 | 0.006 | 5.40E+05 | 6.85E+03 | 3.98E+00 | 0.1 | FALSE | Uncultured bacterium FL0428B_PF12needseq (FJ716468), 98.6% | *Ca*. Magnetobacterium bavaricum TM-1 (LACI01000304), 89.8% | *Thermoclostridium caenicola* EBR596 (AB221372), 83.6% | *Ca*. Magnetobacterium bavaricum TM-1 (LACI01000304), 89.8% | *Nitrospirota* |
| DUR021 | 0.009 | 0.022 | 1.961 | 9.08E+03 | 1.66E+03 | 1.30E+03 | 81.3 | TRUE | Uncultured NKB19 bacterium QEDV3AH03 (CU920011), 97.4% | *Ca*. Hydrogenedentes bacterium MJ-time_bin-1056 (CAIYET010000682), 96.7% | *Wenzhouxiangella marina* KCTC 42284 (CP012154), 84.2% | *Ca*. Hydrogenedentes bacterium MAG_17971_hgd_130 (DUZN01000164), 92.3% | *Hydrogenedentota* |
| DUR022 | 1.775 | 0.274 | 0.023 | 1.79E+06 | 2.06E+04 | 1.53E+01 | 0.1 | FALSE | Uncultured bacterium AN231 (GQ859848), 99.1% | *Geobacter* sp. RPFA-12G-4 (LC379581), 89.5% | *Geobacter soli* GSS01 (JXBL01000001), 88.8% | *Ca*. Belliniella magnetica LBB04 (MK632188), 87.3% | *Thermodesulfobacteriota* |
| DUR023 | 0.341 | 0.356 | 0.042 | 3.44E+05 | 2.68E+04 | 2.79E+01 | 0.1 | FALSE | Uncultured bacterium 450cmOC0043 (KC922900), 99.1% | *Povalibacter uvarum* Zumi 37 (AB548216), 88.8% | *Povalibacter uvarum* Zumi 37 (AB548216), 88.8% | Magnetococcales bacterium WMHbin3 (PDZX01000011), 81.6% | *Pseudomonadota* |
| DUR024 | 0.182 | 2.726 | 0.053 | 1.84E+05 | 2.05E+05 | 3.51E+01 | 0.0 | FALSE | Uncultured bacterium B0618R001_D14 (AB658268), 98.7% | *Ignavibacterium album* JCM 16511(CP003418), 85.8% | *Ignavibacterium album* JCM 16511(CP003418), 85.8% | *Ca*. Magnetomicrobium cryptolimnococcus XYC (JAGYWI010000050), 78.8% | Unclassified Bacteria |
| DUR025 | 0.351 | 0.320 | 0.107 | 3.54E+05 | 2.41E+04 | 7.10E+01 | 0.3 | FALSE | Uncultured bacterium clone 4_mid (MF942713), 99.8% | Proteobacteria bacterium IMCC26096 (MW884648), 99.1% | *Nitrosospira tenuis* Nv1 (FOBH01000006), 93.8% | *Desulfamplus* sp. nJC1bin9 (JADFZQ010000075), 88.8% | *Pseudomonadota* |

**Supplementary Table S4.** Statistics of genomes reconstructed in this study.

| **Parameter** | **DUR002** | **DUR003** |
|---|---|---|
| GenBank assembly accession no. | JAJAPY000000000 | JAJAPZ000000000 |
| Completeness (%) | 94.38 | 75.84 |
| Contamination (%) | 0 | 3.37 |
| GTDB classification | | |
| Phylum | *Elusimicrobiota* | *Elusimicrobiota* |
| Class | *Endomicrobia* | *Elusimicrobia* |
| Order | *Endomicrobiales* | F11 |
| Family | - | F11 |
| Genus | - | FEN-1177 |
| Species | - | - |
| No. of scaffolds | 29 | 137 |
| Assembly $N_{50}$ (bp) | 185704 | 46489 |
| Genome coverage (×) | 218.6x | 79.5x |
| Assembly size (bp) | 3394123 | 2891406 |
| GC content (%) | 39.76 | 52.84 |
| No. of genes | 2714 | 2426 |
| No. of rRNA genes | 3 | 3 |
| 5S rRNA | 1 | 1 |
| 16S rRNA | 1 | 1 |
| 23S rRNA | 1 | 1 |
| No. of tRNA genes | 45 | 49 |
| MAG quality | High-quality | Medium-quality |
